# Microstructural Brain Correlates of Inter-individual Differences in Respiratory Interoception

**DOI:** 10.1101/2024.04.08.588519

**Authors:** Niia Nikolova, Jesper Fischer Ehmsen, Leah Banellis, Malthe Brændholt, Melina Vejlø, Francesca Fardo, Micah Allen

## Abstract

Interoception, the perception and integration of physiological signals, is a fundamental aspect of self-awareness and homeostasis. While previous work has explored interoceptive processing in relation to the cardiac system, research in the respiratory domain, particularly in relation to brain structure and function, is limited. To address this gap, we utilised a Bayesian psychophysical model to quantify perceptual, metacognitive, and affective dimensions of respiratory interoception in a sample of 207 healthy participants. We also measured individual whole-brain microstructural indices of myelination, myeloarchitecture, and cortical iron using quantitative brain imaging. Voxel-based quantification analyses revealed distinct patterns of cortical microstructure in the insular, cingulate, and primary sensory cortices, which underpin interoceptive perceptual sensitivity and precision. In addition, metacognitive bias was associated with increased myelination of the cingulate cortex and periaqueductal grey, while metacognitive sensitivity correlated with myelination of the midline prefrontal cortex. At an affective level, sensitivity to respiratory resistance was related to the myelination of the primary somatosensory cortex. By revealing specific histological brain patterns tied to individual differences in respiratory interoception, our results uncover the neural pathways that govern perceptual, metacognitive, and emotional facets of interoceptive processing.

## Introduction

Interoception, the sensing and processing of internal bodily signals, is critical for higher cognitive processes such as emotion and self-awareness (Berntson & Khalsa, 2021; Craig, 2003; Seth, 2013). Interoception is critical for the regulation of internal states and the maintenance of allostasis (i.e., proactive self-regulation), and has been shown to play a critical function in the regulation of emotion, consciousness, and decision-making (Nikolova, Waade, et al., 2021). Respiratory interoception (hereafter *respiroception*) stands out among the interoceptive modalities owing to its relative ease of manipulability and its association with respiratory and psychiatric conditions such as asthma (Dahme et al., 1996; Harrison, Marlow, et al., 2021), chronic obstructive pulmonary disease (COPD) (Giardino et al., 2010), as well as anxiety and panic disorders (Harrison, Köchli, et al., 2021; Paulus, 2013). To uncover the neural basis of respiroception, we utilised recent developments in quantitative neuroimaging to inter-relate *in vivo* markers of cortical histology to individual differences in perceptual, metacognitive and affective dimensions of interoception.

Interoceptive ability is typically described using a dimensional model which differentiates objective perceptual sensitivity to physiological signals from metacognitive sensitivity (i.e., interoceptive accuracy and awareness, respectively) (Garfinkel et al., 2015, 2016). Metacognition in this context refers to the accurate monitoring of perceptual or cognitive processes, and is quantified by the correlation of subjective confidence (i.e., interoceptive sensibility) with decision accuracy (Fleming & Lau, 2014). Recent computational work suggests that respiratory interoception can be further decomposed through a predictive processing lens (Allen et al., 2022; Brændholt et al., 2023). This framework emphasises a hierarchical view in which sensory prediction errors, which signal for example changes in respiratory resistance or effort, are gated by their precision, or inverse uncertainty. Alterations in interoceptive precision are thought to drive individual differences in perceptual, metacognitive, and affective levels (Ainley et al., 2016; Nikolova, Waade, et al., 2021), and in the cardiac domain has been linked to a range of psychiatric illnesses (Smith et al., 2020, 2021). While a recent study linked respiroceptive precision to the activation of the insula (Harrison, Köchli, et al., 2021), the importance of inter-individual variability in respiroception across these dimensions and hierarchical levels has not been thoroughly investigated.

The psychophysiology underpinning the sensation and perception of breathing is complex and involves both interoceptive and exteroceptive signals. Throughout the respiratory cycle, airflow through the mouth, nose, and airways into the lungs is detected by thermosensory and tactile receptors. Meanwhile, chemoreceptors signal variations in arterial blood gases (Nishino, 2011), and the physical expansion and contraction of the lungs activate stretch receptors distributed around the chest cavity and diaphragm. This rhythmic information is relayed to key brainstem and midbrain nuclei in the medulla and pons, such as the nucleus tractus solitarii and periaqueductal grey as well as subcortical structures including the thalamus and amygdala. From there, respiroceptive brain pathways convey afferent signals to the somatosensory cortex (Pattinson et al., 2009; Raux et al., 2007), and onward to higher-order networks (Davenport & Vovk, 2009; Schroijen et al., 2020; von Leupoldt et al., 2008), primarily involving the medial prefrontal (Biskamp et al., 2017), insular (Davenport & Vovk, 2009), and anterior cingulate cortices (von Leupoldt et al., 2008).

While the functional pathways supporting respiroception are relatively well established, the contribution of variation in the structural configuration of these regions has not been thoroughly examined. Inter-individual differences in brain microstructure, indexing the myelination and/or neurobiological integrity of cortical units, could reflect variation in the function of these pathways and therefore underpin individual respiroceptive abilities (Kanai et al., 2010). Recent advances in quantitative MR imaging (Callaghan et al., 2014; Weiskopf et al., 2015) allow for the mapping of contrasts that are sensitive to histological properties such as myeloarchitecture, iron, and macromolecule content (e.g., myelination, oligodendrocytes and other support structures). To investigate the neuroanatomical basis of individual respiroceptive abilities, we employed voxel-based quantification (Draganski et al., 2011) to correlate brain microstructure with individual interoceptive profiles spanning perceptual, metacognitive and affective levels of processing.

## Materials and Methods

### Participants

A total of 565 (360 females, 205 males) participants (median age = 24, age range = 18 - 56) took part in the Visceral Mind Project, a large-scale neuroimaging project at the Center of Functionally Integrative Neuroscience, Aarhus University. Participants were recruited via the SONA participation pool system and through local advertisements such as posters and social media. The inclusion criteria required participants to have corrected-to-normal vision and fluency in Danish or English. Participants did not take any medications except contraceptives or over-the-counter antihistamines. Furthermore, participants were compatible with standard MRI scanning requirements (i.e., no metal implants, claustrophobia, and not pregnant/breastfeeding). The participants from this dataset took part in multiple tasks, MRI scans, physiological recordings, as well as psychiatric and lifestyle inventories, spread over three visits occurring on different days. The MRI and respiratory interoception data reported in the present study were collected on different days, and the participants were compensated for their participation. The local Region Midtjylland Ethics Committee granted ethical approval for the study and all participants completed written informed consent. The study was conducted in accordance with the Declaration of Helsinki.

### Respiratory Interoception Task

Respiratory interoceptive ability was assessed by the Respiratory Resistance Sensitivity Task (RRST, Nikolova, Harrison, et al., 2021). This task uses Psi, a Bayesian psychophysical procedure to adaptively estimate subject-level thresholds and precision (Kontsevich & Tyler, 1999) for the detection of inspiratory resistances. We further assessed metacognitive bias through average confidence ratings, as well as metacognitive performance, which reflects the trial-level correspondence between confidence ratings and performance accuracy. Participants were asked to breathe through a respiratory circuit, which is automatically compressed by a custom device with fine granularity to produce varying resistance loads (**Figure 1 a**.). In a two-interval forced choice (2IFC) design, each trial encompassed two inhalations and a decision of which of the two was more difficult. This was followed by a visual analog scale (VAS) confidence rating scale of the confidence that the response on that trial was correct, on a range from “Guess” to “Certain” (**Figure 1 b**.). Participants performed a total of 120 trials over the course of approximately 35 minutes, with > 2min breaks at intervals of 20 trials.

**Figure 1.**
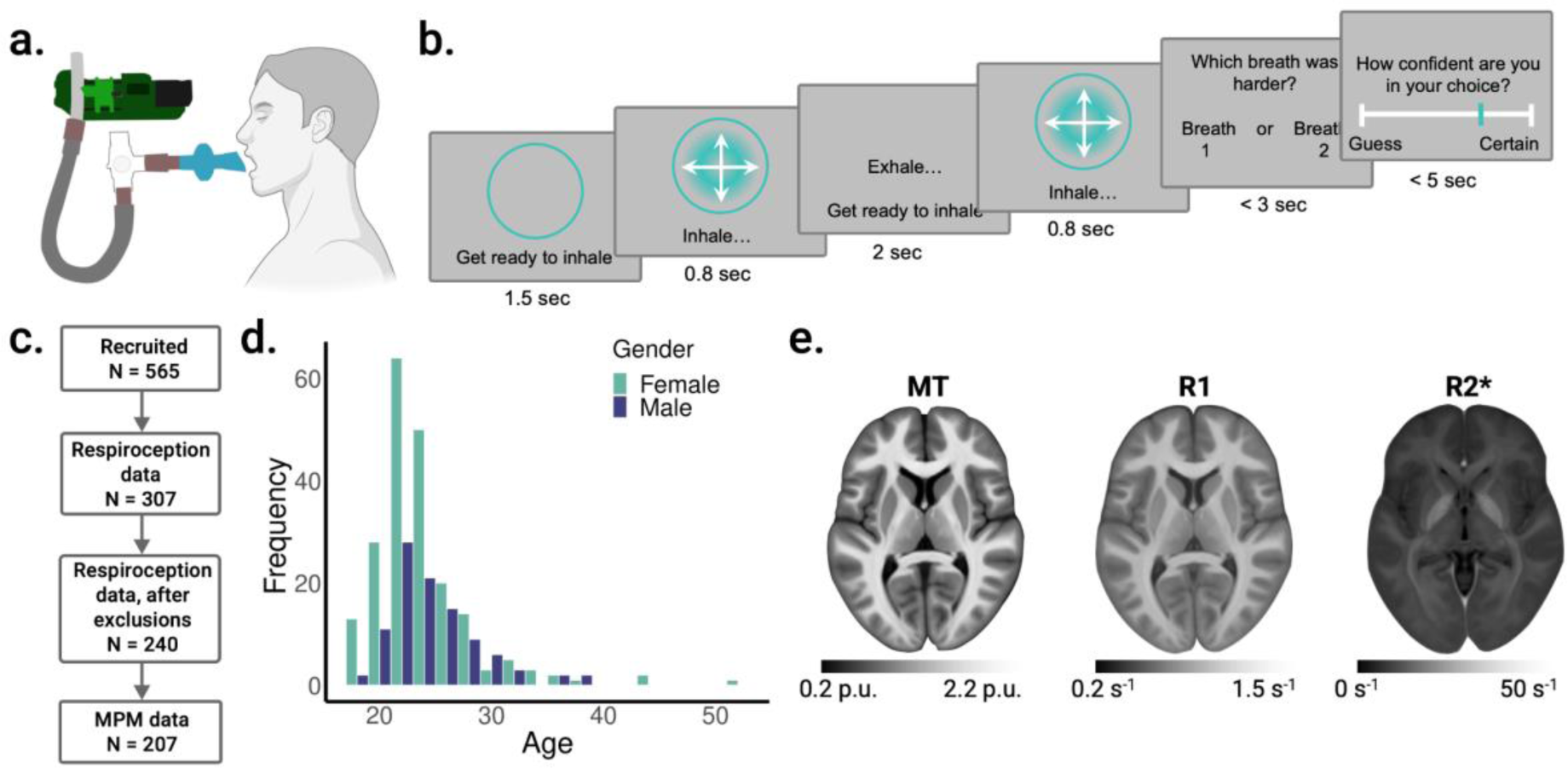
Respiratory resistance discrimination task (RRST) and dataset overview. **a.** Schematic representation of the task apparatus and experimental set-up. Participants breathe through a respiratory circuit on which titrated resistive loads are delivered via the stimulus computer. **b.** Trial schematic depicting the 2IFC design of the task. On each trial participants view a circular cue instructing them to prepare to inhale. The circle then blinks and begins expanding, with the participant instructed to sharply inhale with the expansion of the circle. The participant then exhales and takes a second breath, and a resistive load is randomly applied on either breath one or two. This procedure of pacing the participant’s breathing via visual cues is a novel feature of the RRST, and is intended to reduce intra- and inter-subject variance in respiratory effort. Following the two breaths, the participant indicates by keyboard press whether the first or second breath was more difficult. Across trials, the difference between the loaded breaths is controlled via psychophysical staircasing (see Methods). **c.** Participant inclusion and exclusion flow chart, representing the rigorous data quality controls that were applied to behavioural and neuroimaging data. **d.** Distributions of age and gender of the final sample of 207 participants included in the study. **e.** Average tissue microstructure maps from 442 participants for Magnetization transfer (MT), longitudinal relaxation rate (R1), and transverse relaxation rate (R2*). The scale bars represent the estimated physical values of tissue properties in each map, quantified in standardised units, and represent percent units (p.u.) for MT maps, and per second (s^-1^) for R1 and R2* maps.

We modelled participant RRST behaviour using a Bayesian psychophysical model which fitted a Weibull psychometric function with a guess rate γ of 0.5 and a lapse rate λ of 0.02 (**Eq. 1**) to the trial-level responses, for each participant independently. For this model, trials were coded as correct if participants accurately identified the interval containing a resistance load. The estimated threshold corresponds to the perceptual sensitivity (interoceptive accuracy) or the estimate of parameter α in **Eq. 1**, and the slope, or signal uncertainty (precision), is the value of the β parameter. All analyses of individual differences in respiratory interoception utilised the Psi-estimated psychophysical parameters.

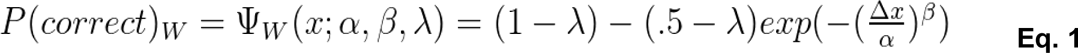

To examine metacognitive performance, we applied a signal theoretic model to distinguish metacognitive sensitivity (i.e., the sensitivity of confidence ratings to objective accuracy) from metacognitive bias (i.e., the overall tendency to report low or high confidence) (Fleming & Lau, 2014). In the context of interoception, metacognitive bias is sometimes called “interoceptive sensibility” (Garfinkel et al., 2016).

Metacognitive sensitivity was estimated by calculating the area under the type 2 (i.e., metacognition) receiver operating characteristic (aROC) curve, a non-parametric measure which indexes the relationship between metacognitive “hits” and “misses” (Fleming et al Science) (Fleming et al., 2010). Metacognitive bias was calculated as the average confidence rating across all trials, irrespective of accuracy. For the estimation of aROC, the continuous confidence ratings were binned into four equally spaced intervals (Galvin et al., 2003; Maniscalco & Lau, 2012). This binning strategy, based on guidelines established in previous studies (Fleming, 2017; Fleming & Lau, 2014; Maniscalco & Lau, 2012), normalises the confidence quantiles for each participant and minimises the risk of empty bins, thereby enhancing the robustness of metacognitive sensitivity estimation.

To probe the affective dimension of respiratory interoception, participants reported subjective levels of ‘unpleasantness’ using a VAS scale ranging from ‘Not unpleasant’ to ‘Very unpleasant’ at intervals of 20 trials throughout the task (i.e., before each break).

### Statistical Analysis

#### RRST Variables

To evaluate the correlations between estimated RRST variables included in the analysis, we calculated Spearman coefficients (see **Supplementary figure 2**). A repeated measures ANOVA was performed to assess the effect of sampling time point on displeasure ratings, with age and gender as covariates. The means and standard deviations of the displeasure ratings are presented in **Supplementary table 1**. As Mauchly’s test of sphericity indicated that the sphericity assumption was violated, χ^2^ (14) = 230.74, the degrees of freedom were corrected using Greenhouse-Geisser sphericity estimates (ε = 0.42). These statistical analyses were performed in JASP (0.18.3) (*JASP*, 2024).

#### Hierarchical Modelling of Respiroceptive Psychophysics

To estimate the overall psychophysical function at a population level (**Figure 3 a**.), we applied a post-hoc hierarchical Bayesian model (Gelman & Hill, 2006; McGlothlin & Viele, 2018). This comprehensive model integrated data from all trials and participants, employing a Weibull function for the fit. The model was characterised by fixed parameters for both the guess rate (establishing the lower asymptote) at 0.5 and the lapse rate at 0.05. Alpha (i.e., threshold) and beta (i.e. slope) were estimated as free parameters.

#### Multi-Parameter Brain Mapping

We used a well-established qMRI protocol (Weiskopf et al., 2013, 2015) to map percent saturation due to magnetization transfer (MT), longitudinal relaxation rate (R1) and effective transverse relaxation rate (R2*), followed by voxel based quantification to relate individual differences in respiroception to patterns of brain microstructure.

### Data Acquisition

The imaging data were collected using a 3T MR system (Magnetom Prisma, Siemens healthcare, Erlangen, Germany), using a standard 32-channel radiofrequency (RF) head coil and a body coil. A set of high-resolution whole brain T1-weighted anatomical images (0.8 mm^3^ isotropic) were acquired using an MP-RAGE sequence (repetition time = 2.2 s, echo time = 2.51 ms, matrix size = 256 × 256 × 192 voxels, flip angle = 8°, AP acquisition direction). Whole-brain image acquisitions at isotropic 0.8 mm resolution were obtained using an MPM quantitative imaging protocol (Callaghan et al., 2019; Weiskopf et al., 2013). The sequences consisted of three spoiled multi-echo 3D fast low angle shot (FLASH) acquisitions and three additional calibration sequences in order to correct for RF transmit field inhomogeneities. The FLASH sequences were acquired with MT, PD and T1 weighting. A flip angle of 6° was used for the MT- and PD-weighted images, and 21° for the T1-weighted acquisitions. MT-weighting used a Gaussian RF pulse 2 kHz off resonance with 4 ms duration and a nominal flip rate of 220°. The field of view was 256 mm head-foot, 224mm anterior-posterior, and 179mm right-left. Gradient echoes with alternating readout gradient polarity were acquired using equidistant echo times ranging from 2.34 to 13.8ms (MT) or 18.4ms (PD and T1), using a readout bandwidth of 490 Hz/pixel. For the MT-weighted acquisition, only 6 echoes were collected to achieve a repetition time (TR) of 25ms for all FLASH volumes. For accelerated data acquisition, partially parallel imaging was performed using the GRAPPA algorithm, with an acceleration factor of 2 in each phase encoded direction and 40 integrated reference lines. A slab rotation of 30° was used for all acquisitions. The B1 mapping acquisition comprised 11 measurements with the nominal flip rate ranging from 115° to 65° in 5° steps. The total scanning time for the qMRI acquisitions was approximately 26 minutes.

### Map Creation

All qMRI images were preprocessed using the hMRI toolbox v. 0.5.0 (January 2023) (Tabelow et al., 2019) and SMP12 (version 12.r7771, Wellcome Trust Centre for Neuroimaging, http://www.fil.ion.ucl.ac.uk/spm/), to correct the raw qMRI images for spatial transmit, receive field inhomogeneities and obtain quantitative MT, PD, R1 and R2* estimate maps. Apart from the enabling of imperfect spoiling correction, the hMRI toolbox was configured using the standard settings. All images were reoriented to MNI standard space prior to map creation. This processing produced four maps modelling different aspects of tissue microstructure: an MT map sensitive to myeloarchitectural integrity (Helms et al., 2008), a PD map representing tissue water content, and R1 map reflecting myelination, iron concentration and water content (primarily driven by myelination) (Lutti et al., 2014), and an R2* map sensitive to tissue iron concentration (Langkammer et al., 2010).

The unified segmentation approach (Ashburner & Friston, 2005) was used to segment MT saturation maps into grey matter (GM), white matter (WM) and cerebrospinal fluid (CSF) probability maps. Tissue probability maps based on multi-parametric maps developed by Lorio et al. (Lorio et al., 2016) were used, without bias field correction given that MT maps do not show significant bias field modulation. The GM and WM probability maps were then used to perform inter-subject registration using Diffeomorphic Image Registration (DARTEL), a nonlinear diffeomorphic algorithm (Ashburner, 2007). The MT, PD, R1 and R2* maps were then normalised to MNI space (at isotropic 1 mm resolution) using the resulting DARTEL template and participant-specific deformation fields. The nonlinear registration of the quantitative maps was based on the MT maps due to their high contrast in subcortical structures, and a WM-GM contrast in the cortex similar to T1 weighted images (Helms et al., 2009). Finally, tissue-weighted smoothing was applied using a kernel of 4 mm full width at half maximum (FWHM) using the voxel-based quantification (VBQ) approach (Draganski et al., 2011). Note that, in contrast to voxel based morphometry analysis, this VBQ smoothing approach aims to minimise partial volume effects and optimally preserves the quantitative values of the original qMRI images by not modulating the parameter maps to account for volume changes. The resulting GM segments for each map were used for all statistical analyses. For visualisation purposes, an average MT map in standard space was generated based on all 442 participants in the study (**Figure 1 e**.).

### Participant Exclusion Criteria and MRI Quality Control

Following inspection, several participants were removed from all analyses for reasons related to either MRI or behavioural data. Three participants were excluded immediately following MR data collection for medical reasons (one cerebral palsy and two other suspected brain abnormalities). Multi-parameter mapping contrast images were acquired for 503 total participants. Three participants were excluded due to errors in the scanning sequences. As MPM data is known to be particularly sensitive to motion artefacts, a thorough quality control (QC) assessment was performed to identify high-motion images. Visual QC HTML reports were created of each participant using the hMRI-vQC toolbox (Sherif et al., 2022), and all reports were visually inspected and labelled by two researchers. Doubtful cases were discussed, and a further 57 participants were excluded due to excessive motion affecting the tissue-class segmentation. Data from the remaining 443 participants was used in the spatial analysis and template creation using DARTEL. Of the remaining participants, 307 completed the RRST. Participants who were behavioural outliers (based on the interquartile range; Q1 - 1.5IQR, Q3+1.5IQR) on the threshold, slope, accuracy or aROC estimates, were excluded from further analyses as these values can indicate a failure of the staircasing procedure and/or a poor understanding of the task instructions. These criteria resulted in excluding 67 participants, leaving 240 participants with RRST data for further analyses. The overlap between the RRST and MPM data left 207 participants for VBQ analyses (145 females, 62 males, median age = 24, age range = 18-52, see **Figure 1 c. & d**.).

### Data Availability

All original code, raw psychophysical data and anonymized summary variables are publicly available on <u>GitHub</u>. Summary statistical brain maps are available on <u>NeuroVault</u>. The raw neuroimaging data reported in this study comprise part of a larger dataset and will be made public as part of a dedicated paper following the passage of a mandatory data embargo which protects participants data privacy rights.

### Voxel Based Quantification Analysis

Grey and white matter masks were generated based on our samples, by averaging the smoothed, modulated GM and WM segment images, and thresholding the result at p > 0.2. Inter-subject variation in MT, R1 and R2* GM maps were modelled in separate multiple linear regression analyses. The RRST threshold, slope, mean confidence, aROC, and mean displeasure were used as regressors of interest in the VBQ analysis (see **Supplementary table 2** for a full list of regressors included). We further controlled for variables known to influence respiratory physiology and/or interoception, modelling daily cigarette smoking, self-reported tendency for intranasal versus intraoral breathing, as well as respiratory interoceptive sensibility as measured by items #4, 11, 21, 26 on the multidimensional assessment of interoceptive awareness (MAIA-II; Mehling et al., 2018). Finally, we included age, gender, body mass index (BMI) and total intracranial volume (TIV) as nuisance covariates in all analyses, following recommended procedures for computational neuroanatomy (Ridgway et al., 2008). All positive and negative main effects analyses were small-volume corrected within the GM mask. Whole-brain maps of each positive and negative t-contrast were analysed using a cluster corrected FWE-cluster p-value with p < 0.001 inclusion threshold (Hupé, 2015; Ridgway et al., 2008). All statistical analyses were conducted in SPM12, and the JuBrain Anatomy Toolbox v. 3.0 (Eickhoff et al., 2005) was used to determine anatomical labels and regional percentages. See **Figure 2** for the MPM analysis pipeline.

**Figure 2.**
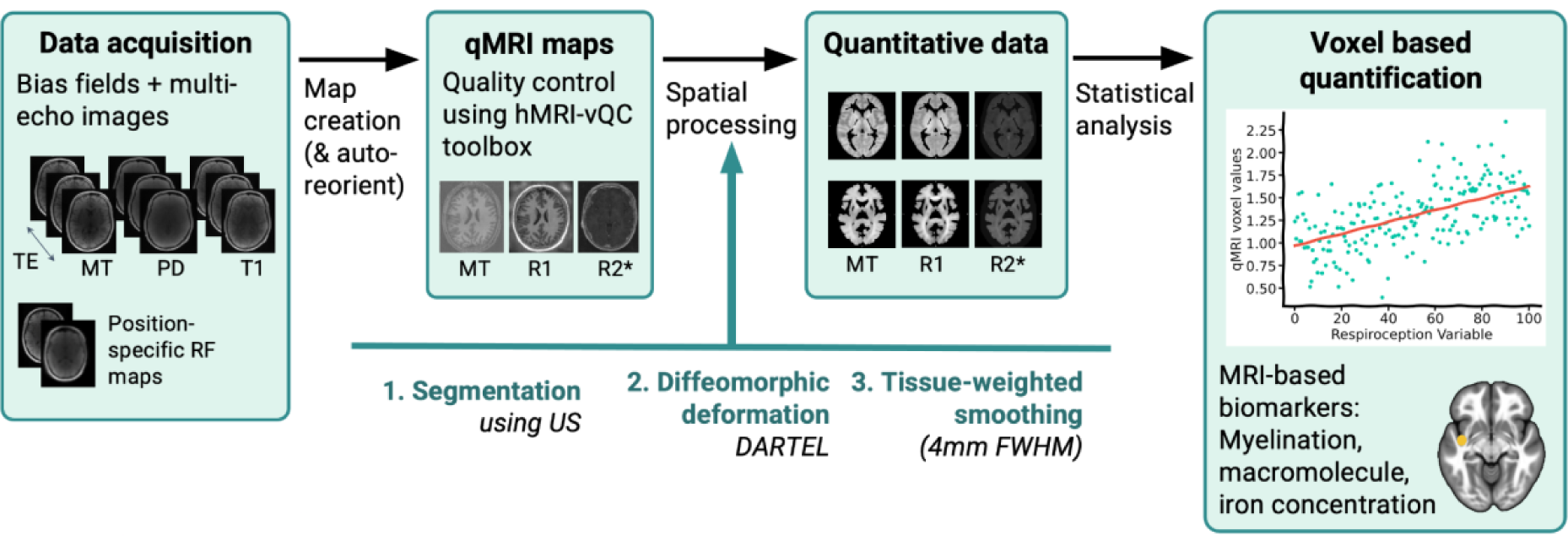
Multi-parameter mapping analysis pipeline. Multi-echo MPM images and bias fields were acquired for MT, PD and T1 contrasts, alongside position-specific RF maps. Auto-reorientation and map creation using the hMRI toolbox produced qMRI maps for MT, R1 and R2*. A combination of automated and manual quality control was performed on these maps, followed by spatial processing including segmentation, diffeomorphic deformation (DARTEL) and tissue-weighted smoothing. The resulting quantitative images were analysed using voxel based quantification in a multiple linear regression analysis, to identify locations in each map type that relate to respiratory interoception parameters while controlling for various nuisance covariates.

## Results

### Respiroception Psychophysics

#### Perception

We first analysed behavioural respiratory interoception data from the RRST, using the hierarchical bayesian fit to the trial-wise obstruction percentage. Across our sample, the mean threshold was 72.70% obstruction (mean, s.d. = 11.86, range: 26.08 - 98.75). The average slope, corresponding to the sensory precision, was 3.44 (mean, s.d. = 3.13, range: 0.10 - 6.53, **Figure 3 a**.). Note that for the Weibull function, a lower slope estimate corresponds to greater response precision. The Psi method effectively controlled the session-level performance at a rate of approximately 70 - 80% correct responses (**Figure 4 d**.), assuring that further analysis of metacognition was not confounded by differences in accuracy performance.

**Figure 3.**
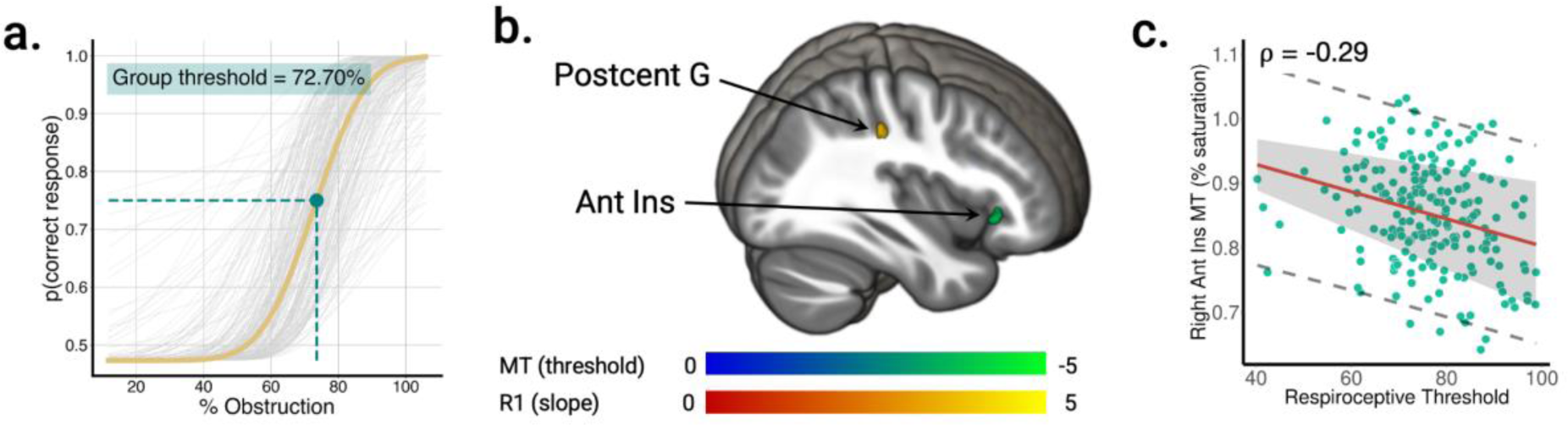
RRST psychophysics and VBQ results for the perceptual level. **a.** Grand mean psychometric fit (yellow) overlaid on individual PMF fits (grey), demonstrating that average respiratory thresholds are approximately 70% airway obstruction (teal point), with substantive inter-individual variance around this value. PMF fits are based on hierarchical estimates. **b.** and **c.** R1 map values in the postcentral gyrus are positively correlated to respiroceptive precision (slope parameter), while MT map values in the anterior insula correlate negatively to respiroceptive accuracy (threshold). Maps are FWE-cluster corrected for multiple comparisons at pFWE < 0.05.

**Figure 4.**
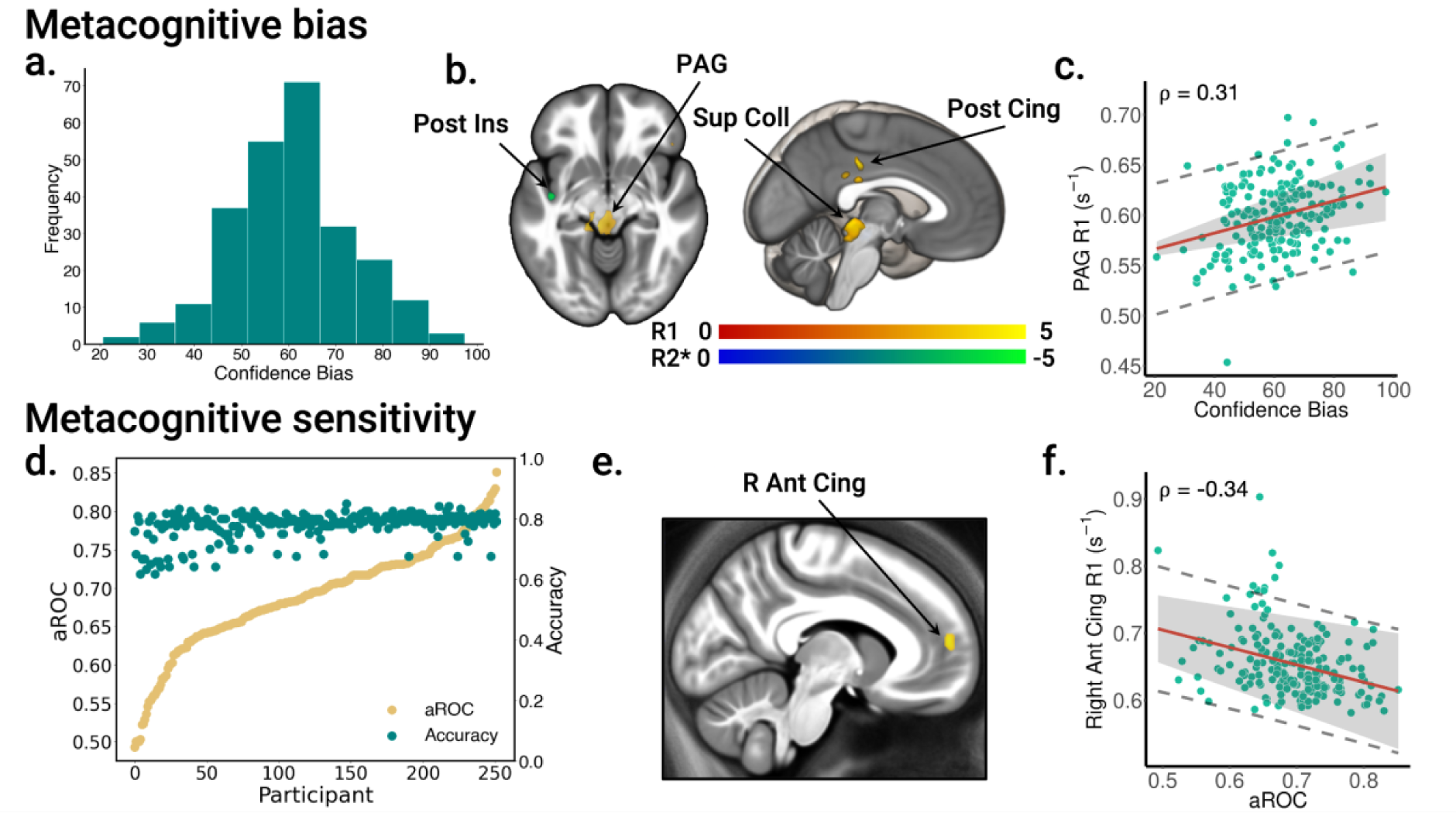
Brain microstructural correlates of metacognitive bias and sensitivity. **a.** Distribution of mean confidence ratings across the sample. **b.** Positive (red - yellow) and negative (blue - green) confidence bias R1 map contrast. **c.** R1 values in the Superior Colliculus and Periaqueductal Grey correlate positively with confidence bias. **d.** Accuracy (teal) and metacognitive sensitivity (aROC, yellow), sorted by each participant’s aROC estimate. Participants show substantial variations in metacognition while accuracy is held relatively constant by the Psi staircase procedure. **e.** R1 values in the anterior cingulate cortex are negatively correlated to metacognitive sensitivity (aROC). **f.** R1 values in the right Anterior Cingulate Cortex correlate negatively with metacognitive sensitivity as indexed by aROC. Maps are FWE-cluster corrected for multiple comparisons at pFWE < 0.05.

#### Metacognition and Affective Ratings

With respect to the metacognition parameters, we estimated the confidence bias, or the tendency to report low or high confidence, by taking the mean of confidence ratings over the entire session for each participant. We observed substantial variability in confidence bias across participants, with a range from 22 - 91% (**Figure 4 a**.). We further calculated the area under the type 2 receiver operating characteristic (aROC) curve, which estimates metacognitive sensitivity while accounting for confidence bias. Whereas perceptual performance was controlled at 70 - 80% accuracy, aROC estimates varied from 0.51 to 0.85 (**Figure 4 d**.).

Ratings of unpleasantness, indexing respiroceptive resistance-induced negative affect, showed that the task was perceived as mildly aversive, with median ratings of 22.92%. However, individual participants showed substantial variability, both in their mean rating (S.D. = 28.16%, 95% C.I. = 26.22 - 33.34), as well as in the pattern of ratings across blocks (**Figure 5 a**.). We replicated our previous results (Nikolova et al., 2022) showing that there is large inter-individual variability in subjective unpleasantness ratings throughout the task. We tested whether there was a significant effect of time within participants using a rmANOVA (factor of timepoint, Greenhouse-Geiser sphericity correction), and found no significant linear effect of timepoint (F(3.200, 611.108) = 2.086, p = .097).

**Figure 5.**
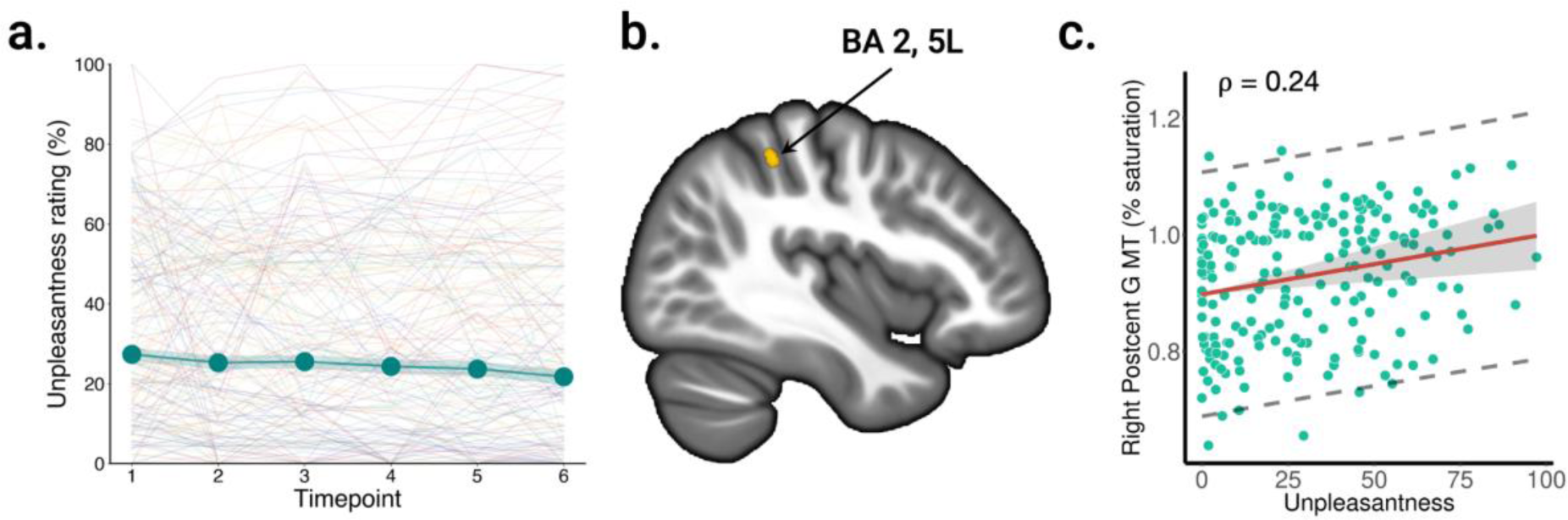
Brain microstructural correlates of affective ratings. **a.** VAS scale ratings of how ‘unpleasant’ participants found the RRST, collected at intervals of 20 trials. Thin lines represent ratings of individual subjects, while teal markers show median ratings at each timepoint and shaded area represents the standard error. While the mean of these affective ratings shows minimal variability over the course of the session, there is inter-individual variability in the affective scores. **b.** and **c.** Mean affective ratings are positively correlated with MT map values in the right postcentral gyrus. Maps are FWE-cluster corrected for multiple comparisons at pFWE < 0.05.

To estimate the inter-dependence of perception, metacognition and affect ratings, we calculated Spearman correlation coefficients between each pair of variables. This revealed that respiroceptive threshold was moderately related to slope and unpleasantness ratings (slope: r(250) = 0.20, p < 0.005; unpleasantness: r(250) = 0.17, p < 0.005), whereas the metacognition measures aROC and mean confidence were not correlated to any of the other variables (**Supplementary figure 2 b.**). A benefit of the RRST is that it allows for the estimation of the threshold and slope, enabling their relative contributions to be controlled for and assessed independently in the subsequent VBQ analysis. We also observed that gender was correlated with threshold (r(250) = -0.27, p < 0.005), unpleasantness (r(250) = -0.18, p < 0.005) and BMI (r(250) = 0.22, p < 0.005), and that age correlated negatively with slope (r(250) = -0.20, p < 0.005), and positively with BMI (r(250) = 0.23, p < 0.005).

### Voxel Based Quantification

To examine the neurobiological correlates of inter-individual variability in respiroceptive processing, we related parameter estimates of perceptual variables (threshold, slope), metacognitive variables (mean confidence, aROC), and affect (unpleasantness ratings) to brain microstructure. We found that MT saturation in the right anterior insula (*k* = 314; *p*FWE_corr_ < 0.001; peak voxel coordinates: x = 46 y = 24 z = −3) and left anterior cingulate cortex (*k* = 290; *p*FWE_corr_ < 0.001; peak voxel coordinates: x = 3 y = 54 z = 17) was negatively correlated with respiroceptive threshold (**Supplementary figure 2 b.**), indicating that greater myeloarchitectural integrity was associated with lower respiroceptive sensitivity. Meanwhile, R1 in the right postcentral gyrus (*k* = 289; *p*FWE_corr_ < 0.001; peak voxel coordinates: x = 37 y = -30 z = 39) related positively with PMF slope estimates (**Figure 3b**.). These results suggest that the ability to accurately discriminate respiratory loads, as indicated by a reduced threshold, is associated with myeloarchitecture in the anterior portions of the cingulate cortex and insula. The precision in detecting resistive loads is instead related to local myelination in primary somatosensory cortex, so that reduced myelination confers greater precision.

Individual variability in confidence bias, or the tendency to provide high or low confidence ratings regardless of performance, was positively correlated with R1 map values in the right posterior cingulate cortex (*k* = 408; *p*FWE_corr_ < 0.001; peak voxel coordinates: x = -3 y = 28 z = 38) and left superior parietal cortex (*k* = 792; *p*FWE_corr_ < 0.001; peak voxel coordinates: x = 33 y = -52 z = 65). Interestingly, we also observed associations with the R1 values in the midbrain, namely the periaqueductal grey and superior colliculus (*k* = 5260; *p*FWE_corr_ < 0.001; peak voxel coordinates: x = -13 y = -30 z = −13); **Figure 4 b**.), indicating that greater myelination in these regions was associated with a positive confidence bias. The R2* values in the left posterior insula meanwhile, were negatively correlated to metacognitive bias, suggesting that high iron concentrations in this region was associated with overall lower mean confidence ratings (*k* = 211; *p*FWE_corr_ = 0.014; peak voxel coordinates: x = 46 y = 24 z = −3); **Figure 4 b**.).

While metacognitive bias relates an individual’s propensity to be over- or under-confident in their performance on a task, their metacognitive sensitivity indexes how well their confidence ratings relate to the trial-by-trial accuracy. R1 map values in the left anterior cingulate cortex (*k* = 866; *p*FWE_corr_ < 0.001; peak voxel coordinates: x = -11 y = 53 z = 11) and frontal pole (*k* = 191; *p*FWE_corr_ = 0.01; peak voxel coordinates: x = -41 y = 46 z = -4) were negatively related to metacognitive sensitivity, as indexed by the type 2 aROC, indicating the involvement of this region in metacognition and performance monitoring.

We found that variability in mean unpleasantness ratings induced by respiratory resistance correlated positively with MT in areas BA 2 and 5L of the postcentral gyrus (*k* = 271; *p*FWE_corr_ < 0.001; peak voxel coordinates: x = -40 y = -40 z = 50; **Figure 5 b., c**.), suggesting that a greater myeloarchitectural integrity may indicate higher affective sensitivity to the inspiratory resistive loads, rendering them more aversive.

**Table 1:** Summary of whole-brain VBQ results.

| Whole brain Results |  |  |  |  |  |  |  |  |
| --- | --- | --- | --- | --- | --- | --- | --- | --- |
| Contrast | Map | Region | k | p(FWE-corr) | Z value | x | y | z |
| Threshold - | MT | R Anterior Insula | 314 | < 0.001 | 4.68 | 46 | 24 | -3 |
|  |  | L Anterior Cingulate | 290 | < 0.001 | 4.62 | 3 | 54 | 17 |
| Slope + | R1 | L Postcentral Gyrus | 289 | < 0.001 | 4.22 | 37 | -30 | 39 |
| Mean Confidence + | R1 | Midbrain (PAG) | 5,260 | < 0.001 | 4.74 | -13 | -30 | -13 |
|  |  | L Superior Parietal | 792 | < 0.001 | 5.06 | 33 | -52 | 65 |
|  |  | R Posterior Cingulate | 408 | < 0.001 | 4.53 | -3 | 28 | 38 |
| Mean Confidence - | R2* | L Posterior Insula | 211 | 0.014 | 4.21 | -41 | -10 | -9 |
| aROC - | R1 | L Anterior Cingulate | 866 | < 0.001 | 5.38 | -11 | 53 | 11 |
|  |  | L Frontal Pole | 191 | 0.01 | 3.92 | -41 | 46 | -4 |
| Mean Displeasure + | MT | L Postcentral Gyrus | 271 | < 0.001 | 3.93 | -39 | -40 | 50 |

## Discussion

This study investigated the microstructural brain correlates of respiratory interoception across perceptual, metacognitive and affective levels. For this purpose, we employed a Bayesian psychophysical method to measure respiroceptive ability alongside quantitative MRI profiles in a sample of over 200 participants. The present work represents the largest investigation of respiroception to date, and also the first brain imaging study to utilise the respiratory resistance sensitivity task (RRST), which differentiates interoceptive sensitivity and precision. We replicated previous results indicating a population level threshold resistance sensitivity of approximately 70% obstruction, and found similar levels of inter-individual variation in both sensitivity and precision parameters (Nikolova et al., 2022). Here, we exploited this variability to understand how each dimension of respiroception maps onto local differences in brain microstructure.

### Microstructural Brain Underpinnings of Respiroceptive Perception

We showed that the myeloarchitectural integrity of the anterior insular cortex (AIC) was related to reduced respiroceptive thresholds and thus greater sensitivity to resistive loads. Greater myeloarchitectural integrity in the AIC therefore may confer an improved ability to discriminate inspiratory resistance. Previous studies have thoroughly established the involvement of the AIC in processing visceral information (Barrett & Simmons, 2015; Craig, 2003; Seth, 2013) and breathlessness (Faull & Pattinson, 2017). Interestingly, while the posterior insular cortex is sometimes referred to as “primary viscerosensory cortex” (Craig, 2002; Critchley & Harrison, 2013; Nieuwenhuys, 2012), the AIC is known to have a more multi-modal profile. This is due in part to the unique agranular cytoarchitecture and connectivity of the region, which has long-distance projections across both cortical and subcortical structures and connects to limbic, sensory and prefrontal areas (Menon & Uddin, 2010; Sridharan et al., 2008). These connections allow the AIC to be involved in integrating sensory stimuli, extracting salience cues, and communicating multimodal inputs in a modulatory way across the brain (Allen, 2020; Seth, 2013). As discriminating resistive loads inevitably requires the integration of various interoceptive and exteroceptive sensory cues, this finding highlights the integrative nature of the region in determining respiroceptive sensitivity.

Respiroceptive precision, as measured by the slope of the psychometric function, was associated with myelination in the primary somatosensory regions of the postcentral gyrus that correspond to the mouth and throat areas (Belyk & Brown, 2014; Haggard & de Boer, 2014; Miyamoto et al., 2006). This connection echoes previous findings that the structure of primary sensory areas is important for individual visual acuity and sensitivity (Duncan & Boynton, 2003; Schwarzkopf et al., 2011). Interestingly, our study highlights that respiratory sensitivity and precision are tied to unique microstructural characteristics in somatosensory and viscerosensory regions, suggesting respiroception uniquely recruits different branches of the body-related neural pathways. Neurobiologically, post-mortem studies indicate that myelination strongly correlates with R1 contrast in somatosensory areas (Lutti et al., 2014), while magnetization transfer (MT) is more reflective of macromolecular content, indicating a possible role for glial and support cells. Thus, respiroceptive precision might be linked to myelination in the primary somatosensory cortex’s grey matter, while interoceptive sensitivity could depend more on the integrity and connectivity of the brain’s myeloarchitecture, particularly within the anterior insular cortex (AIC). Future studies utilising combined functional and structural brain imaging will be needed to understand the contributions of these microstructural features to respiratory processing.

### Microstructural Brain Underpinnings of Respiroceptive Metacognition and Affect

Our findings further showed a relationship between metacognitive processing and cortical myelination. Specifically, respiratory interoceptive sensibility (i.e., mean confidence or metacognitive bias), correlated with myelination in distributed cortical regions such as the posterior parietal and posterior cingulate cortices. In the exteroceptive domain, previous studies found neural correlates of subjective confidence in similar brain regions (Bang & Fleming, 2018; Kiani & Shadlen, 2009), suggesting that there may be multimodal confidence representations spanning interoceptive and exteroceptive modalities in these areas. In contrast, exteroceptive metacognitive sensitivity was previously linked to the activity, myelination, and volume of the rostrolateral prefrontal cortex (Allen et al., 2017; Fleming et al., 2010, 2012), whereas here we found that the myelination of the ventromedial prefrontal cortices (VMPFC) correlated with respiroceptive metacognitive sensitivity. The VMPC has been implicated in diverse aspects of brain-body interaction, self-processing, and interoception, and this finding may therefore highlight more modality specific aspects of metacognition for breathing (Azzalini et al., 2019; D’Argembeau, 2013; Nagai et al., 2004).

In addition to these cortical findings, we also showed that myelination levels in midbrain regions including the periaqueductal grey (PAG) and superior colliculus were positively related to confidence bias. The PAG is a core structure in the central autonomic network (Benarroch, 1993; Berntson & Khalsa, 2021; Saper, 2002), critical to neuromodulation in the perception of dyspnea, as well as homeostatic regulation, affective responses, nociception and stress (Bandler et al., 2000; Faull et al., 2019; Faull & Pattinson, 2017; Sclocco et al., 2018). Increased myelination in the PAG could therefore contribute to the salience of ascending respiratory signals, leading to increased interoceptive sensibility for breathing.

Finally, subjective affective ratings related positively to myeloarchitecture in the postcentral gyrus and intraparietal sulcus, in a region more dorsal to the effect we observed for interoceptive precision. This region of the somatosensory cortex has been shown to be involved in attentional regulation and anxious arousal (Brown et al., 2023), and it has been proposed that neuromodulation of the nearby intraparietal sulcus (IPS; e.g., using TMS) could be used to reduce attention to threat and anxious arousal in clinical samples. The IPS exhibits strong functional connectivity within the fronto-parietal attention network (Cole et al., 2014), as well as with subcortical regions such as the locus coeruleus, known to be involved in anxiety (Liebe et al., 2022). Our findings therefore suggest a relationship between affective state and myeloarchitecture in somatosensory cortex, and carry potential implications for attentional regulation and anxious arousal.

### Future Directions and Limitations

An advantage of our methodology is the inherent interpretability of quantitative MRI (qMRI) units, facilitating direct comparisons with future investigations into the neural substrates of interoceptive processing, and providing a normative sample of respiroceptive values and their cortical correlates in brain microstructure. Our findings also raise questions about how genetic and environmental factors shape local brain microstructure. Previous research highlights the impact of life experiences on cortical grey matter volume, with studies in both rodents and humans showing that early-life dietary iron changes predicting alterations in cortical myelination, iron levels, and cognitive function (Greminger et al., 2014; Radlowski & Johnson, 2013). Exploring factors such as nutrition, stress, and adversity further, particularly through prospective designs (e.g., Karcher & Barch, 2021), could offer valuable insights into how early-life experiences influence the neural circuits supporting respiratory interoception.

Our study extends previous work linking exteroceptive perceptual abilities and variation in cortical structure to the domain of respiratory interoception (Schwarzkopf et al., 2011). However, while our results provide new information about the relationship between brain structure and respiroception, they cannot speak to the underlying causal mechanisms linking brain function, structure, and behaviour. Our findings thus motivate future work aimed at unravelling how the microstructural brain variability revealed here might underpin specific respiratory computational mechanisms, and also could motivate prospective cohort research seeking to understand the aetiology of these structure-function relationships.

One possible drawback of our study is the lack of physiological measures (i.e., mouth pressure and airway resistance) when estimating respiroceptive psychophysics. The correlation between tube obstruction and airway resistance follows a nonlinear equation, such that under constant airflow static resistance remains fixed for a given level of obstruction in the RRST. Nevertheless, the force exerted during inhalation can alter air flow in the compressed area (the segment of silicone tubing in the breathing circuit), affecting differential pressure. By measuring the air circuit’s mechanical characteristics, such as airflow and pressure, one could calculate the static resistance generated by a stimulus. This approach would allow for adjusting stimulus threshold values based on physiological parameters unique to each trial, thereby eliminating potential biases due to variations in individual breathing patterns, like the force and duration of inhalation. Although we continuously recorded airflow and differential pressure during the task, a technical issue resulted in a loss of stimulus triggers, preventing analysis of this data. Nonetheless, our previous validation study demonstrated that the relationship between tube obstruction and both resistance and pressure follows a consistent log-linear pattern across different participants and stimulus levels (Nikolova et al., 2022). This consistency is due to our task design, which uses visual cues to standardise the force and duration of inspiratory breaths across trials, ensuring reliable threshold estimation even without physiological data.

In conclusion, this study advances our understanding of respiratory interoception by revealing distinct cortical underpinnings associated with interoceptive accuracy, precision, and metacognitive processes. Our findings reveal previously unknown specificity in the cortical and subcortical respiroceptive brain hierarchy, indicating that regionally specific differences in brain microstructure underlie variations in respiratory interoception across multiple functional levels. The respiroceptive brain profiles identified here can aid in the design of more targeted investigations probing the neural mechanisms of respiroceptive ability and metacognition, their clinical relevance, and the relative combinations of genetic factors and prior experiences in shaping interoceptive processing.

## Acknowledgements

NN, LB, MB, MN and MA are supported by a Lundbeckfonden Fellowship (under Grant R272-2017-4345), and a European Research Council Grant (ERC-2020-StG-948788). JFE and FF are supported by a European Research Council Grant (ERC-2020-StG-948838).

## Notes

### Competing Interest Statement

The authors have declared no competing interest.

