## Supplementary material for "Microstructural Brain Correlates of Inter-individual Differences in Respiratory Interoception"

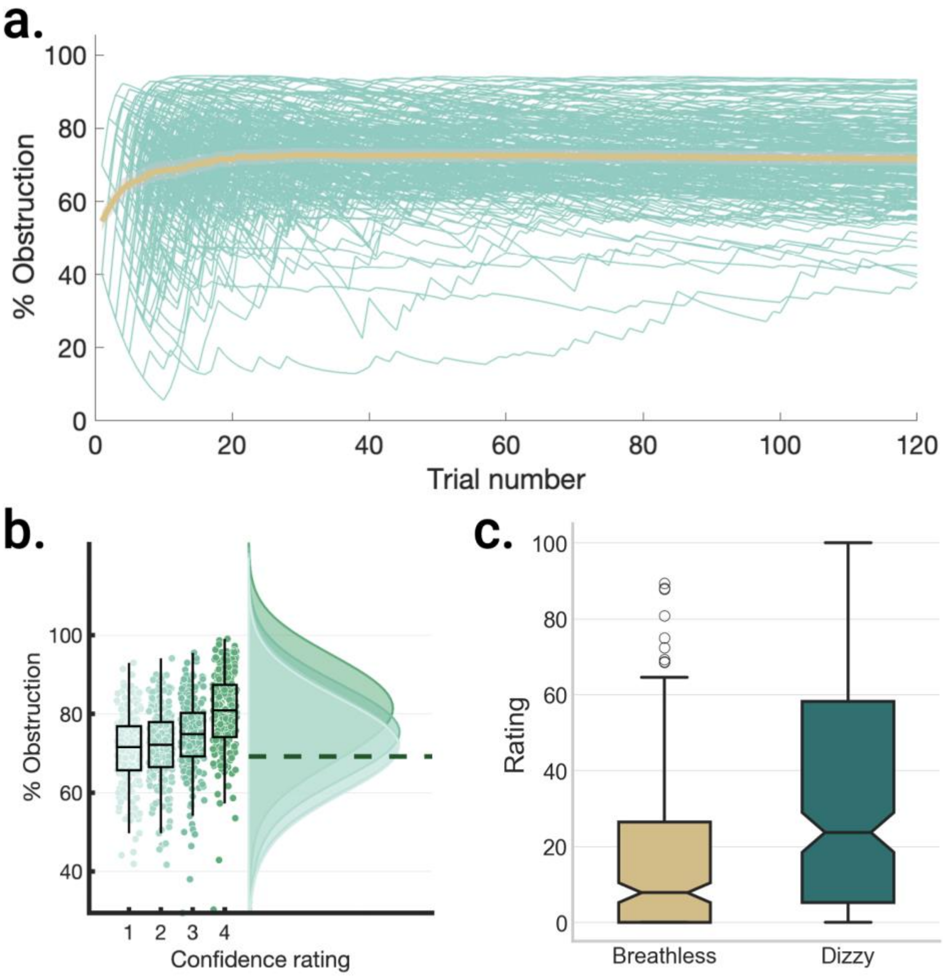


**Supplementary figure 1. a.** Threshold estimate traces for all 207 participants included in the final analysis over 120 trials of an RRST session (teal), and the mean threshold estimate over the sample (yellow). **b.** Stimulus intensity (i.e., percentage obstruction) co- varies with confidence bin input to the metacognition model, showing the subjective correspondence between stimulus intensity and confidence. **c.** Boxplot depicting mean dizziness and breathlessness symptoms across participants. Bar height represents mean ratings, error bars denote SEM, and grey circles show individual participants' ratings.

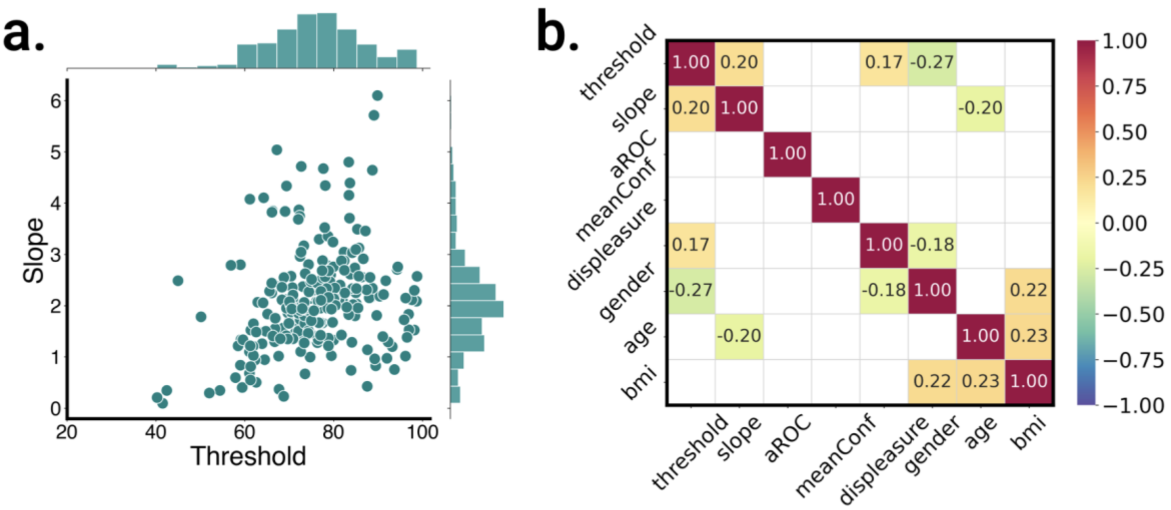


**Supplementary figure 2.**

**a.** Distributions of threshold and slope estimates for all participants. **b.** Correlations between perceptual, metacognitive and affective RRST parameters, and demographic variables. Heatmap shows only significant FDR corrected Spearman correlation coefficients.

### Repeated Measures ANOVA

**Within Subjects Effects**

| Cases | Sphericity Correction | Sum of Squares | df | Mean Square | F | p | $\eta^2_p$ |
|---|---|---|---|---|---|---|---|
| Timepoint | None | 1877.222 [a] | 5.000 [a] | 375.444 [a] | 2.086 [a] | 0.065 [a] | 0.011 |
| | Greenhouse-Geisser | 1877.222 | 3.200 | 586.720 | 2.086 | 0.097 | 0.011 |
| Timepoint ✻ age | None | 2789.668 [a] | 5.000 [a] | 557.934 [a] | 3.099 [a] | 0.009 [a] | 0.016 |
| | Greenhouse-Geisser | 2789.668 | 3.200 | 871.902 | 3.099 | 0.024 | 0.016 |
| Timepoint ✻ gender | None | 1811.157 [a] | 5.000 [a] | 362.231 [a] | 2.012 [a] | 0.075 [a] | 0.010 |
| | Greenhouse-Geisser | 1811.157 | 3.200 | 566.072 | 2.012 | 0.107 | 0.010 |
| Residuals | None | 171919.049 | 955.000 | 180.020 | | | |
| | Greenhouse-Geisser | 171919.049 | 611.108 | 281.324 | | | |

*Note.* Type III Sum of Squares

[a] Mauchly's test of sphericity indicates that the assumption of sphericity is violated (p < .05).

**Between Subjects Effects**

| Cases | Sum of Squares | df | Mean Square | F | p |
|---|---|---|---|---|---|
| age | 1012.134 | 1 | 1012.134 | 0.284 | 0.595 |
| gender | 26870.389 | 1 | 26870.389 | 7.547 | 0.007 |
| Residuals | 679994.045 | 191 | 3560.178 | | |

*Note.* Type III Sum of Squares

*Assumption Checks*

**Test of Sphericity**

| | Mauchly's W | Approx. $X^2$ | df | p-value | Greenhouse-Geisser ε | Huynh-Feldt ε | Lower Bound ε |
|---|---|---|---|---|---|---|---|
| Timepoint | 0.295 | 230.739 | 14 | < .001 | 0.640 | 0.652 | 0.200 |

**Supplementary table 1.** Repeated measures ANOVA of displeasure ratings over timepoints, including age and gender as covariates. As the sphericity assumption was violated, and epsilon ε < 0.75, the Greenhouse-Geisser sphericity correction was used.

| Variable | Abbreviation | Category |
|---|---|---|
| Threshold (psi) | psi_thresh | Respiroceptive accuracy |
| Slope (psi) | psi_slope | Respiroceptive precision |
| aROC | aROC | Metacognitive sensitivity |
| Mean confidence | meanConf | Metacognitive bias |
| Mean displeasure | meanDisp | Affect |
| Smoking amount | | Control covariates |
| Breathe nose/mouth | | Control covariates |
| MAIA 4 | | Control covariates |
| MAIA 11 | | Control covariates |
| MAIA 21 | | Control covariates |
| MAIA 26 | | Control covariates |
| Gender | | Nuisance covariates |
| Age | | Nuisance covariates |
| BMI (body mass index) | | Nuisance covariates |
| TIV (total intracranial volume) | | Nuisance covariates |

**Supplementary table 2**. Full list of regressors included in the VBQ analysis. Threshold and slope estimates from the Psi model, aROC, mean confidence and mean displeasure were included as regressors of interest, along with a number of control covariates known to influence respiratory physiology and interoception. Abbreviations: aROC, (type 2) area under the receiver operating characteristic curve. MAIA, multidimensional assessment of interoceptive awareness. BMI, body mass index. TIV, total intracranial volume.
